# Translating surveillance data into incidence estimates

**DOI:** 10.1101/443952

**Authors:** Yoann Bourhis, Timothy R. Gottwald, Frank van den Bosch

## Abstract

Monitoring a population for a disease requires the hosts to be sampled and tested for the pathogen. This results in sampling series from which to estimate the disease incidence, *i.e*. the proportion of hosts infected. Existing estimation methods assume that disease incidence is not changing between monitoring rounds, resulting in underestimation of the disease incidence. In this paper we develop an incidence estimation model accounting for epidemic growth with monitoring rounds sampling varying incidence. We also show how to accommodate the asymptomatic period characteristic to most diseases. For practical use, we produce an approximation of the model, which is subsequently shown accurate for relevant epidemic and sampling parameters. Both the approximation and the full model are applied to stochastic spatial simulations of epidemics. The results prove their consistency for a very wide range of situations.

## Introduction

Monitoring programs are used to keep track of the invasion and spread of human, animal and plant pathogens. They are often structured in discrete rounds of inspection, during which subsamples of the host population are assessed for disease status (Parnell et al., 2017). Given a sequence of monitoring rounds, a key question in interpreting these data is the estimation of the incidence^1^ of the disease in the host population. There are two special cases of this general question that have received some attention.

Firstly, monitoring is often motivated by the need for early responses to enable eradication or containment. For example, *early detection* of the disease permits reduced cullings of animal and plant hosts (Carpenter et al., 2011; Cunniffe et al., 2015, 2016), as well as reduced resorts to emergency quarantines or travel restrictions for human hosts (applied *e.g*. for SARS, Smith, 2006).

Secondly, monitoring is frequently motivated by the desire of *proving disease absence* from a host population (Caporale et al., 2012), which is of key importance for the transport and trade of hosts. The main question then concerns the sufficient sample size (Cannon, 2002). An example of this is the practical “rule of three” (Louis, 1981; Hanley & Lippman-Hand, 1983). It gives the upper bound of the 95% confidence interval (CI) of the incidence when all of the *N* sampled hosts are assessed as healthy: *q*_95_ = 3/(*N* + 1). Estimating a disease incidence (noted *q* hereafter), or proving its absence, is mostly interesting during the early stages of epidemics, *i.e*. when incidences are low and containment measures are still promising.

Simple practices like the “rule of three” make the assumption that the samples are independent binomial draws of probability *q*. However, epidemics are structured processes, and samples are very likely to carry dependencies to those structures. For example, by pooling all the samples together, we neglect the fact that early monitoring rounds have most likely sampled a lower incidence *q* than the current one, resulting in an underestimation of the incidence. An alternative and unbiased solution consists in estimating *q* only from the last round to date. But obviously, such a poor use of data would only be tolerable in cases where the monitoring interval and epidemic growth rate are both very large, so that the previous monitoring rounds can be deemed uninformative of the last one. The temporal dependence of samples has been addressed by Metz et al. (1983) in the design of appropriate monitoring programs, as well as by Bourhis et al. (2018) for the incidence estimation problem in the specific case of *disease absence, i.e*. when all samples return healthy.

Making use of all monitoring data, we propose here a generalised solution to the incidence estimation problem. Building on the simple logistic equation, we develop an estimation model that accounts for the evolution of the disease during the monitoring period. Following the idea of the rule of three, and in the way of Parnell et al. (2012) and Alonso Chavez et al. (2016), we produce an approximation of this model. Its derivation only requires simple algebraic operations which makes it more suitable for practitioners. The full model and its approximation are shown accurate when tested against stochastic sampling of logistic epidemic simulations. Finally, they are taken one step further and conclusively tested against spatially explicit stochastic simulation models.

## Material and Methods

Monitoring a population for a disease results in sampling series like Table 1. We define *K* as the number of monitoring rounds iterated in time. *N*_*k*_ is the sampling size of monitoring round *k, i.e*. the number of hosts whose pathological status is assessed at time *t*_*k*_. *M*_*k*_ is the number infected hosts detected during round *k*. Finally, Δ_*k*_ is the time interval between monitoring rounds *k* and *k* + 1.

**Table 1.** Variables and structure of a sampling series.

| Monitoring round | 1 | 2 | ... | k | ... | K-1 | K |
| --- | --- | --- | --- | --- | --- | --- | --- |
| Number of samples | $N_1$ | $N_2$ | ... | $N_k$ | ... | $N_{K-1}$ | $N_K$ |
| Number of positives | $M_1$ | $M_2$ | ... | $M_k$ | ... | $M_{K-1}$ | $M_K$ |
| Time interval | $\Delta_1$ | $\Delta_2$ | ... | $\Delta_k$ | ... | $\Delta_{K-1}$ | — |

### One monitoring round

Considering *q* the disease incidence in the population, the probability of any sampling size *N* and respective result *M*, is given by the binomial probability density function

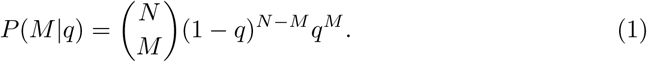

A more general form, accounting for the occurrences of false positives and negatives in the detection process, would be

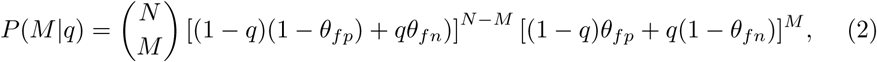

where *θ*_*ƒ n*_ and *θ*_*ƒ p*_ are respectively the rates of false negatives and false positives. But we will not expand this further here.

In a practical context, *q* is the variable that we want to estimate from samples characterised by their size *N* and their outcome *M*. To this end we use Bayes’ rule:

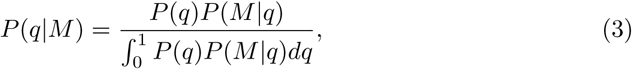

where *P*(*q*|*M*) is the probability density of *q* given *M* and *N*. Assuming no information on the incidence before sampling, we set a uniform prior *P*(*q*), simply resulting in *P*(*q*|*M*) ∝ *P*(*M*|*q*) (Gelman et al., 2003).

### K monitoring rounds

To account properly for the dynamic incidence between monitoring rounds, our proposition is to inform the binomial probability density with an epidemiological component, noted *Z*_*k*_:

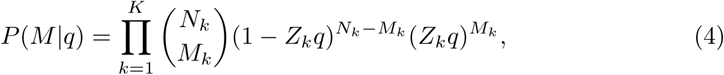

where *M* on the left-hand side represents the whole sampling series, *i.e M*_1_, *M*_2_,…,*M*_*K*_. In Eq. 4, the parameter *Z*_*k*_ ∊ [0,1] modulates the value of *q* for the samples to be compared to the disease incidence that was actually found in the population when they were made (*i.e*. at time *t*_*k*_). For the last monitoring round, *Z*_*k=K*_ = 1, and then decreases with *k* < *K*.

We assume that the disease incidence, *q*, evolves logistically (van der Plank, 1963; Murray, 2002) in time *t* as:

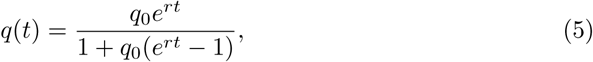

where *q*_0_ is the incidence at time *t*_0_ and *r* is the epidemic growth rate. To include this logistic growth into the binomial probability density, we define *Z*_*k*_ as:

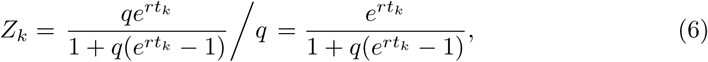

where 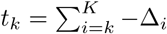 are the sampling dates, with the last date defined as *t*_*K*_ = 0. Eq. 5 can be fed negative time values to derive incidence backward in time (so that *q*_0_ in Eq. 5 is in fact the incidence at the end of monitoring, *i.e*. the one to estimate).

Similarly to the case of one monitoring round, we use Bayes’ rule to get the un-normalised posterior distribution *P*(*q|M*). Practically, it is given by Eq. 4, which is computed for a discretised array of *q* ∊ [0,1], and from which quantiles can be derived (a method called grid approximation, see *e.g*. Kruschke, 2014). This estimation model has been deployed as an online app (see Supplementary Materials for details).

### A useful approximation

As mentioned in the introduction, the upper bound of the confidence interval (CI) is a useful measure of the highest, still likely, incidence we can expect in the population given the outcome of our monitoring program. Deriving an approximation from the estimation model previously described proved itself intractable. However, various methods exist for approximating the CI of a binomial probability density (Wallis, 2013), and they appeared to fit the binomial-shaped probability density given by Eq. 4. After preliminary testing of those methods, we choose the Agresti-Coull interval for its accuracy for low incidences (Agresti & Coull, 1998). The Agresti-Coull interval is defined as

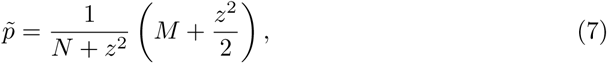

and then

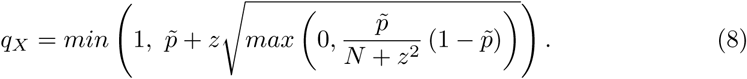

where *q*_*x*_ is the upper limit of the *X*% CI and *z* is the corresponding 1 — *α*/2 quantile of the standard normal distribution. For the one-sided 95% CI that we used in the examples hereafter, *z* = 1.645.

As previously, the estimation of *q*_*X*_ needs to account for the epidemic growth. Because of the density dependence of the logistic equation, we cannot ground this new 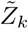 on the logistic model, as it would need *q* to estimate *q*. Therefore, we assume an exponential growth of the disease in the population. In practice, this assumption is realistic as, during early infection, the epidemic growth is exponential, even according to the logistic model (van der Plank, 1963). Then, 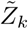 quantifies the disease evolution between rounds as

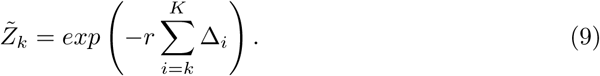

Finally, we aggregate the samples together with respect to the epidemic growth via 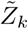:

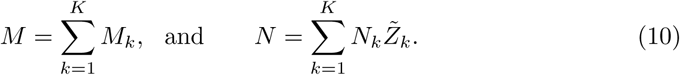

These aggregated values of *M* and *N* are then substituted in Eqs. 7 and 8 to derive *q*_*X*_. By scaling the size of the historic samples with the disease incidence they actually sampled, we adjust their contribution to the total sampling effort. The *min* and *max* operators in Eq. 8 are added to deal with the possibility of having *N* < *M* for some values of 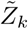.

As discussed in the introduction early detection of epidemics and the establishment of disease absence have received some attention in the epidemiological literature. Two specific approximations have been produced for the estimation of the disease incidence (1) when the first infected hosts are detected (*first discovery* event, Parnell et al., 2012), and (2) while no infected hosts have yet been detected (sampling for *disease absence*, Bourhis et al., 2018). See Supplementary Materials for details. The general estimation model we provide in this study encompasses those specific contexts but extends to any sampling series, being irregularly structured or not, and whatever their outcoming *M*_*k*_

### Asymptomatic period

In most diseases, infected hosts produce symptoms after an asymptomatic period. Often, asymptomatic hosts contribute to the epidemic dynamics by spreading the disease while still undetectable (cryptic) when sampled. The logistic equation handles this period, noted *σ*, as

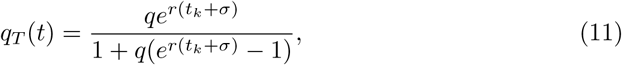

where *q*_*T*_ is the total incidence of the disease, while *q* becomes the detectable incidence (in our problem, *q* is the sampled incidence and *q*_*T*_ the estimated one). Hence, Eq. 6 becomes

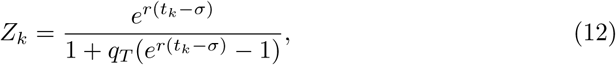

Therefore, *Z*_*k*_ now expresses the ratio *q*(*t*_*k*_)/*q*_*T*_(*t*_*K*_), instead of *q*(*t*_*k*_)/*q*_*t*_(*t*_*K*_), and is then no longer equal to 1 for the last round (if *σ* > 0). For the exponential approximation, the Eq. 9 simply becomes

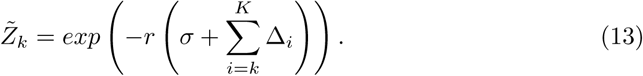

### Testing the model

The consistency of the full model and the accuracy of its approximation are first tested against simulations of stochastic sampling on non-spatial logistic epidemics. We consider a uniform distribution of incidences *q*_*T*_ that we want to estimate individually. For each one of them, a monitoring program is designed with *N*_*k*_ and Δ_*k*_ drawn from Poisson distributions of mean 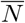 and 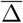. From the logistic equation (Eq. 5), the detectable incidence *q* is derived for every sampling dates *t*_*k*_. Then binomial draws with probability *p* = *q*(*t*_*k*_) and size *n* = *N*_*k*_ simulate the sampling process of the hosts, resulting in *M*_*k*_. For every *q*_*T*_ an exact upper bound of its CI, *q*_*X*_, is derived with the full model, while an approximated one, 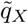 is derived with the approximation. A relevant test then consists in checking that the upper limits of the *X*% CI are above *q*_*T*_ in *X*% of cases. This test is done for contrasted values of the sampling (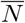 and 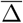) and epidemic parameters (*r* and *σ*).

The full model and its approximation are also tested against a spatial stochastic simulation model. In this case, the epidemics are no longer modelled with the logistic equation but through a transmission rate and a dispersal kernel of the pathogens. To this end, the hosts are distributed in a 2D-space and aggregated randomly in field-like structures mimicking the distribution of the trees in an orchard. Details on this landscape model are given as Supplementary Material. The epidemic progress follows an exponential power kernel (Rieux et al., 2014). The probability of a susceptible individual to become infected in a unit of time is then given by

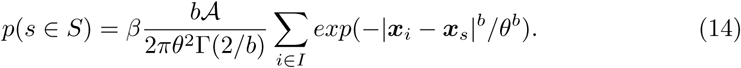

Where *s* is a susceptible host among the set of all susceptible hosts *S*. Similarly, *i* and *I* represent the infected hosts. 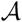 is the area occupied by one host and *Γ* is the Gamma function. *β* is the probability of infection, *θ* is the dispersal scale and *b* is a shape parameter (producing fat-tailed kernels for *b* < 1). The coordinates *x* mark the location of the hosts. Following Klein et al. (2006), the mean dispersal distance for this 2D kernel is given by:

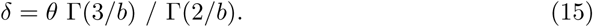

The spatial dynamics are simulated with the *τ*-leap Gillespie algorithm (Keeling & Rohani, 2008). Three sampling methods of increasing realism are tested: random (*i.e*. host locations have no impact on sampling), stratified (*i.e*. sampling equally distributed among fields) and systematic (*i.e*. sampling occurs every *n* hosts along ranks). As no effect of the sampling method is observed, results are shown for the random one. Apart from this, the model and its approximation are evaluated in the same way as the non-spatial case.

## Results

### Model behaviours

Figure 1 illustrates the effects of the epidemic and sampling parameters on the resulting probability densities of the incidence and upper quantiles *q*_95_. Increasing *M*, the number of positive/infected hosts in the sample, unsurprisingly increases the estimated incidence. Increasing the sample size *N* reduces the uncertainty in the estimates. Increasing the sampling interval Δ decreases their impact on the estimation. This reflects the fact that samples taken further back in time are less informative of current disease incidence. Regarding the epidemic parameters, the growth rate *r* and the asymptomatic period *σ* (not shown on Figure 1 for dimensional reasons) have very similar effects to Δ. They both increase the estimated incidence by decreasing the impact of the historic samples, *i.e*. the ones which sample lower incidences *q*. By doing that, Δ, *r* and *σ* reduce the effective sample size (*i.e*. 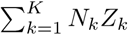), which also contributes in increasing the uncertainty on the estimates (*i.e*. producing densities with larger variance).

**Figure 1.**
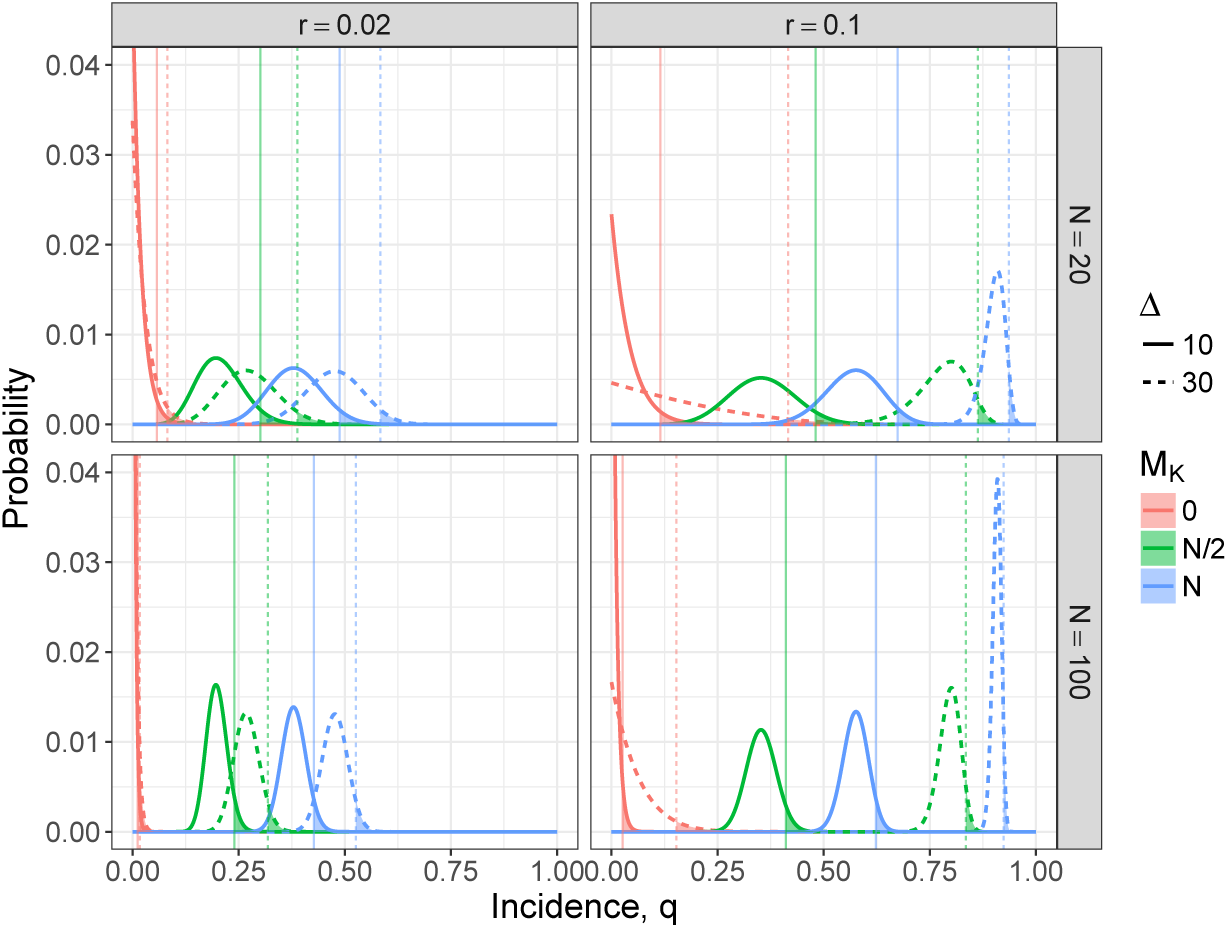
Probability densities of the incidence *q* given by Eqs. 4 and 3. The vertical lines mark the upper limit of the 95% CI. The densities result from a sampling series composed of K=3 monitoring rounds, of which the first two are fully negative (*i.e. M*_1_ = *M*_2_ = 0) and the last varies from *M*_3_ = 0 (*i.e*. all sampled hosts are negative) to *M*_3_ = *N* (*i.e*. all sampled hosts are positive). These probability densities are represented for varying values of epidemic growth rate *r*, sampling size *N* and sampling interval Δ.

### Test against logistic epidemics

Figure 2 shows the distribution of the exact and approximated upper bounds of the 95% CI, q95 and 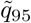, for uniform distributions of *q*_*T*_ and different values of the epidemic and sampling parameters. The full model, which like the simulations builds on the logistic equation, behave exactly as expected: it ensures that 95% of the q95 are above their respective *q*_*T*_, for every set of parameters tested. On the other hand, the approximation displays another behaviour easily explained by its underlying exponential growth model. For the low incidences which are relevant to practice (*i.e*. say *q*_*T*_ < 0.25), the approximation is accurate (the distributions of *q*_95_ and 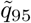 do overlap). For higher incidences, *i.e*. when the logistic growth decelerates unlike the exponential growth, the approximation tends to overestimate the incidence (increasingly with *r*, *σ* and Δ).

**Figure 2.**
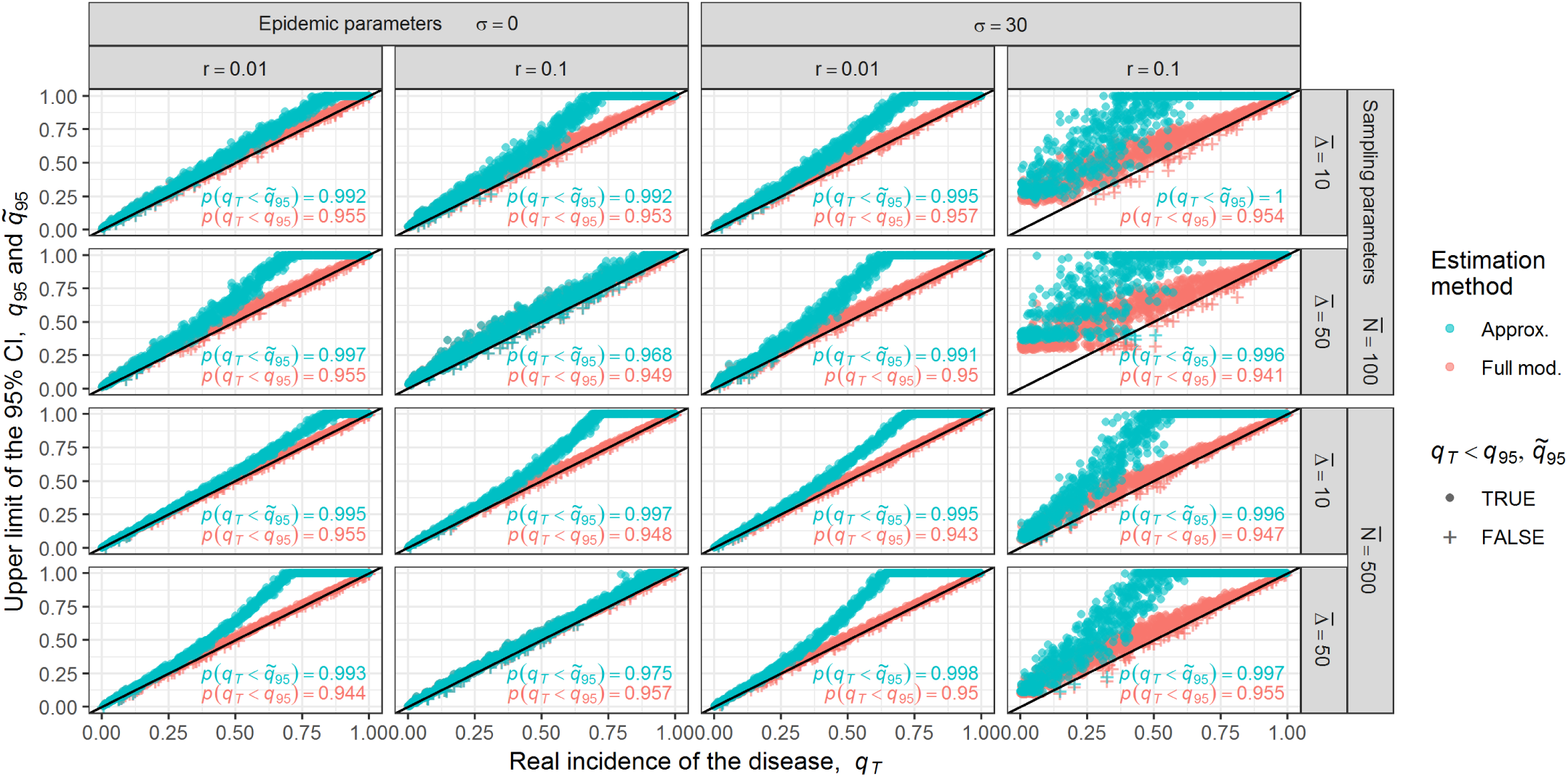
Estimation of *q*_95_ and 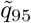 from sampling series of non-spatial epidemics, *i.e*. simulated with the logistic equation (Eq. 5). These estimations are made for contrasted values of sampling and epidemic parameters (and for *K* = 5 monitoring rounds). Using here the 95% CI, we expect 95% of the estimated *q*_95_ and 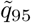 to be above the actual incidence in the field at the end of monitoring *q*_*T*_, i.e. above the oblique black line. The inserted texts summarise these scores for the full model (in red) and its approximation (in blue).

Another model behaviour is of particular interest: when *r* and *σ* are large (*cf*. the rightmost column), we observe that the estimated *q*_95_ and 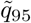 do not align well with the diagonal for small incidences *q*_*T*_. For those cases of very hazardous pathogens with high epidemic growth rates and long asymptomatic periods, the sampling size *N* is too small to allow discrimination between the non-detection cases (*i.e*. the one for which all the *M*_*k*_ = 0), and that larger sampling effort is needed for the estimation to be useful.

Although increasing *r* and *σ* accelerates the divergence between the logistic and the exponential curves, the approximation appears accurate for early infections even considering very high values of epidemic parameters such as *r* = 0.1 day^−1^ or *σ* = 100 days.

### Test against spatial epidemics

When locating the hosts in space, the epidemic becomes driven by two new elements: the dispersal range of the pathogen and the intensity of host clustering (Brown & Bolker, 2004). Both determine how easily the pathogen spreads across the landscape or remains restricted to a local group of hosts. Random distributions of hosts and long dispersal ranges result in smooth progressions of the pathogen across the landscape, following a logistic-like curve. However, as the dispersal range decreases and host aggregation increases, the simulated epidemics will tend to include interruptions between periods of seemingly logistic growth within host clusters. Questions then arise regarding the performance of our estimation model on such epidemics.

In this regard, the estimation model and its approximation are tested for varying host aggregations and dispersal ranges. Host aggregation is summarised by *μ*, the number of hosts in a field (*sensu* host cluster). For a given landscape-scale population of hosts, more hosts by fields means fewer but more populated fields (see the Supplementary Material for an illustration). The dispersal scale *θ* is translated in terms of mean dispersal distance *δ* (see Eq. 15), while *μ* is translated in terms of 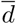, a landscape metric measuring the mean minimal distance between the fields within a landscape (see Euclidean Nearest Distance in Leitao et al., 2006).

In the same way as Figure 2, Figure 3 shows the performance of the model and its approximation for gradients of dispersal scales *θ* (in columns) and host aggregations *μ* (in rows). For each parameter set *θ* and *μ* (*i.e*. each panel in Figure 3), 50 epidemics are first simulated for 50 different landscapes in order to identify the value of *r* producing the best fitting logistic curve. This *r* then informs the incidence estimation model and its approximation for the subsequent testing set of 2000 epidemics and landscapes. Most of the figure is in agreement with expectations: the estimated *q*_95_ do align neatly above the diagonal, showing in practice the accuracy of the estimation model. The approximation appears to be a good simplification of the full model for early detection. However, the estimation model also produces overestimations of the incidence, in bottom row and left column (*i.e*. where the dots do not align above the diagonal), cases for which the distance between host clusters (quantified by 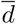) is too large for the pathogen dispersal range (quantified by *δ*), restricting the usefulness of the model to cases where 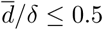.

**Figure 3.**
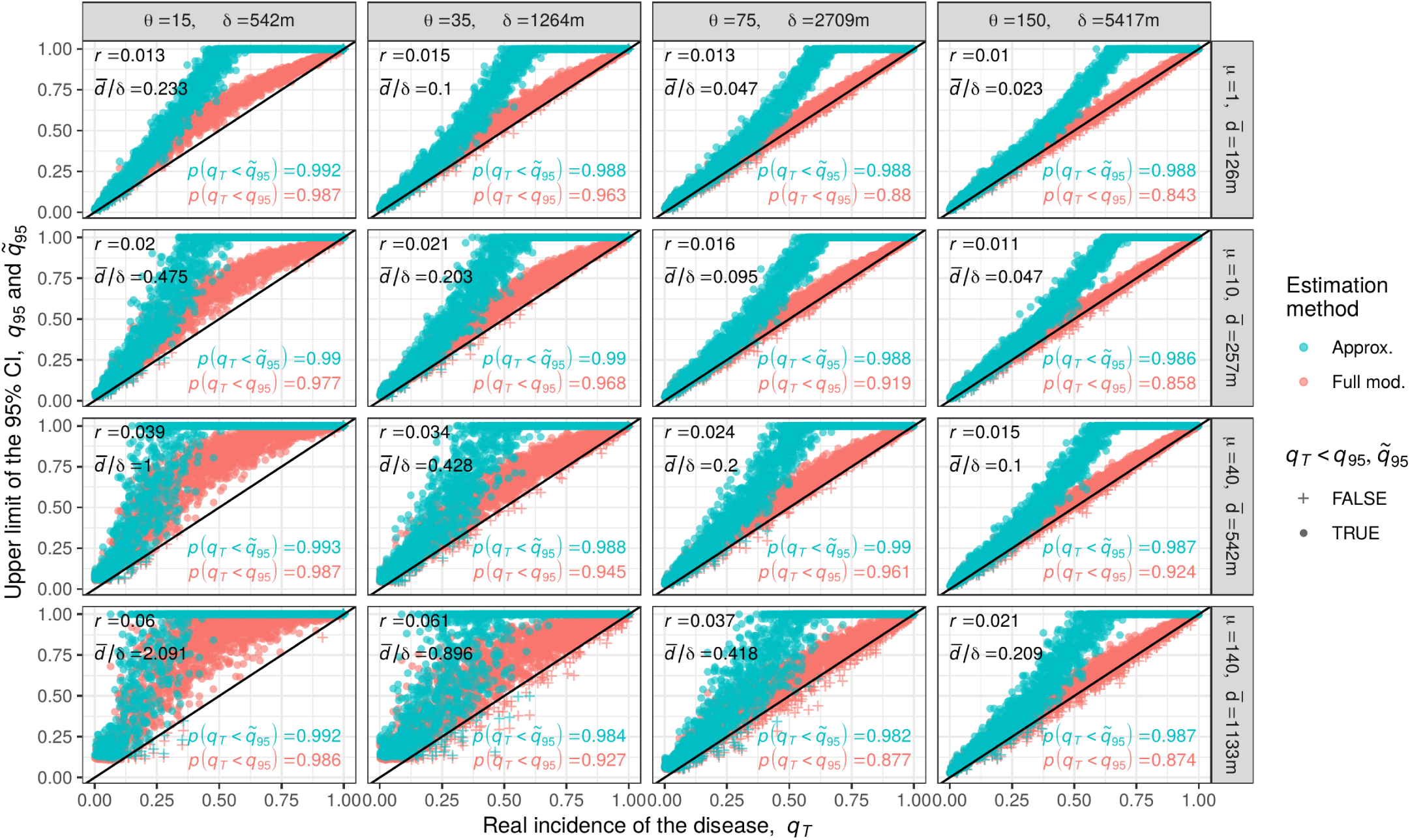
Estimation of *q*_95_ and 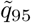 from sampling series realised on spatially explicit epidemics, *i.e*. simulated with the dispersal kernel (Eq. 14). These estimations are made for varying dispersal ranges *θ* and hosts aggregations *μ* while maintaining constant values of the non-spatial parameters (*N* = 100, Δ = 30, *σ* = 30, *K* = 5, as well as *β* = 75 and *b* = 0.45 for the remaining kernel parameters). For better understanding, *θ* and *μ* are shown with their distance translation in meters, *δ* and 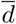. The identified logistic growth rate *r* is given for each experiment. The resulting distributions of *Q*_95_ and 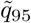 are qualitatively similar for other realistic values of the fixed parameters.

We notice also that *p*(*q*_*T*_ < *q*_95_) can be below the 95% expectation. As stochasticity scatters the realised epidemic curves symmetrically around the fitted logistic one, such effect is mechanical. Yet, this is of no practical concern as, when various epidemic growth estimates are available for a disease, the highest is chosen for caution (and not the mean as we did here).

## Discussion

The model developed in this paper is suitable for many monitoring designs, including those with irregular sampling sizes and time intervals between rounds. The model weights the monitoring outcomes according to an estimation of the population incidence at their respective sampling time, before aggregating them into a single binomial-shaped probability density of the incidence whose quantiles have practical interests. The model is directly applicable for situations in which surveillance is not built on the self-reporting of symptomatic hosts, which makes it appropriate for most animal and plant species.

Deriving the probability density of the incidence from the sampling series is computationally inexpensive, but still requires the use of a computing language. Therefore, we have produced an online app interfacing the full model as exhaustively as possible, as well as an approximation of the model which can be derived with simple algebraic operations. Our intention is to equip the widest audience of practitioners with this incidence estimation capability. The approximation is as flexible as the original model, and we have shown that its inaccuracies are restricted to high level of incidences that are less relevant when dealing with emerging epidemics. However, in case such high incidence estimation is needed, we have seen that the approximation is conservative, i.e. biased towards an overestimation of disease progress.

The model relies on the simple and deterministic logistic equation. That it is consistent with more complex systems is not obvious. The tests presented here against spatial, stochastic and non-logistic based simulations of epidemics, are very promising. They show that our non-spatial model is robust against the decisive impact of spatiality and stochasticity. The model gives accurate estimates of the disease incidence for most simulated epidemics. However, highly aggregated host distributions, as well as short distance dispersing pathogens, support epidemics that diverge from the logistic equation. In those contexts, the disease progression across the landscape is not steady but punctuated by rare events: the pathogen jumps between distant host clusters. Then, the very distinctive trajectories this epidemic can take do not simplify well into a single logistic curve. In such cases, reduced pathogen dispersal and increased host aggregation result in the habitat fragmentation of the pathogen. This allows us to consider the pathogen dispersal between distant host clusters as a primary infection. Theoretically, we consider that the epidemic is composed of multiple smaller epidemics running simultaneously, that can be dealt with individually, or be given multiscale considerations (as in Cameron & Baldock, 1998; Coulston et al., 2008).

Recent technological innovations are changing epidemiological surveillance for more timely and exhaustive censuses. For example, the monitoring of human epidemics is already augmented by the supervision of social networks (Chen et al., 2014) and internet search queries (Yuan et al., 2013; Yang et al., 2015). Tree monitoring could also be assisted by satellite high-resolution imagery (Li et al., 2014; Salgadoe et al., 2018). Those forthcoming innovations will still need robust and epidemiologically informed estimation methods and, even featuring continuous monitoring, there is no reason to see them incompatible with an adaptation of our model. However, in any foreseeable future, most contagions will still be monitored through discrete and censored inspections and hence, remain within the immediate scope of the estimation model presented here.

## Acknowledgements

The work at Rothamsted forms part of the Smart Crop Protection (SCP) strategic programme (BBS/OS/CP/000001) funded through the Biotechnology and Biological Sciences Research Council’s Industrial Strategy Challenge Fund. Authors are also thankful to the Newton Fund of the British Council and the US Department of Agriculture for funding support, as well as to Fran Lopez-Ruiz for its inputs regarding the online app.

## Supplementary Materials

**A** The estimation model as an online app.

**B** Development of the specific approximations for first discovery and disease absence.

**C** Details, illustration and code for the landscape model.

## Footnotes

1 We use here the plant pathology definition where incidence is the fraction of host units infected. In human and other animal pathology this is termed prevalence.

